# Automated tracking of *S. pombe* spindle elongation dynamics

**DOI:** 10.1101/2020.10.09.333765

**Authors:** Ana Sofía M. Uzsoy, Parsa Zareiesfandabadi, Jamie Jennings, Alexander F. Kemper, Mary Williard Elting

## Abstract

The mitotic spindle is a microtubule-based machine that pulls the two identical sets of chromosomes to opposite ends of the cell during cell division. The fission yeast *Schizosaccharomyces pombe* is an important model organism for studying mitosis due to its simple, stereotyped spindle structure and well-established genetic toolset. *S. pombe* spindle length is a useful metric for mitotic progression, but manually tracking spindle ends in each frame to measure spindle length over time is laborious and can limit experimental throughput. We have developed an ImageJ plugin that can automatically track *S. pombe* spindle length over time and replace manual or semi-automated tracking of spindle elongation dynamics. Using an algorithm that detects the principal axis of the spindle and then finds its ends, we reliably track the length and angle of the spindle as the cell divides. The plugin integrates with existing ImageJ features, exports its data for further analysis outside of ImageJ, and does not require any programming by the user. Thus, the plugin provides an accessible tool for quantification of *S. pombe* spindle length that will allow automatic analysis of large microscopy data sets and facilitate screening for effects of cell biological perturbations on mitotic progression.

## Introduction

As the advent of new microscopy techniques has vastly increased the rate of production of cell biological imaging data, computational approaches to tracking image features have become an increasingly important tool for automating the analysis of live microscopy images (1). Analysis pipelines that automate image tracking can decrease time spent analyzing data, supplement the accuracy of the analysis, or allow detection of information that a human observer might miss (2). Furthermore, they have the potential to vastly increase throughput and thereby allow detection of rare image features or of correlations among cells treated with different conditions (3–5).

In the past decade, with the development of several superresolution microscopy techniques (6–11), fast, automatic localization of individual diffraction-limited spots has seen particular advancement (12). Automatic tracking of diffraction-limited spots, such as individual or small clusters of proteins or other molecules (13–15) or randomly-labeled “speckles” that can serve as proxies to identify movement such as sliding or turnover within larger objects (16), have proven very useful for monitoring protein dynamics in live cells (17), including those of the microtubule cytoskeleton (18). These approaches are also important for assembling super-resolution images following the detection and localization of fluorescence from single molecules (19–21).

However, many cellular structures are not diffraction-limited in size or cannot be easily labeled for super-resolution imaging, and detection of those structures requires a distinct approach. Existing tools can help automate the detection and measurement of larger, non-diffraction limited features in cell biological images, such as nuclei and the cellular membrane (22,23), and a novel, open-source Python-based toolkit uses deep learning to segment sub-cellular structures in fluorescence microscopy images (24). Tools for automatic detection of more complex cellular structures and their dynamics are likely to become increasingly important amidst ongoing efforts to create “atlases” that map the landscape of cellular structures and protein-protein interactions (24–26).

One such cellular structure is the mitotic spindle, a microtubule-based machine that segregates the two identical sets of chromosomes to opposite ends of the cell during cell division. Errors in chromosome segregation lead to extra or missing chromosomes, a condition called aneuploidy that is associated with cancer, miscarriage, and birth defects. Furthermore, the assembly and disassembly of the mitotic spindle is an important indicator of the cell cycle, so the spindle can be used both to monitor normal cell cycle progression and to detect defects (27). Tracking the biologically significant changes in spindle shape and position over time thus make tracking of the mitotic spindle a particularly intriguing computational problem. However, few tools for automatic spindle tracking are available. Larson and Bement (28) created a MATLAB package that tracks the rotation angle and pole body location in the mitotic spindle of epithelial cells in *Xenopus laevis* embryos and allowed them to determine the stage of cell division merely by the rotational dynamics of the spindle. Decarreau et al. (29) use MATLAB to semi-automate tracking of the rotation angle of mammalian spindles, but with significant manual input from the user. Both of these existing tools were developed for tracking the spindles of higher eukaryotes, and neither tools are readily applicable to the detection of the spindle of the fission yeast *S. pombe*, which are morphologically quite distinct.

*S. pombe* is an important model organism for studying the mitotic spindle. Its spindle shares many features with higher eukaryotes, while a robust toolkit for genetic manipulation and its relatively simple structure make it easy to perturb (30). The *S. pombe* mitotic spindle consists of a single bundle of microtubules organized by microtubule associated proteins (MAPs), as seen in Figure 1. As mitosis progresses, growing microtubules are slid apart by molecular motors, segregating chromosomes and elongating the spindle. Spindle length in *S. pombe* is quite stereotyped from cell to cell, and is a robust indicator of mitotic progress (31). Thus, measuring spindle elongation dynamics can serve as a proxy to quantify the effects of perturbations to cell division machinery (32–36).

**Figure 1.**
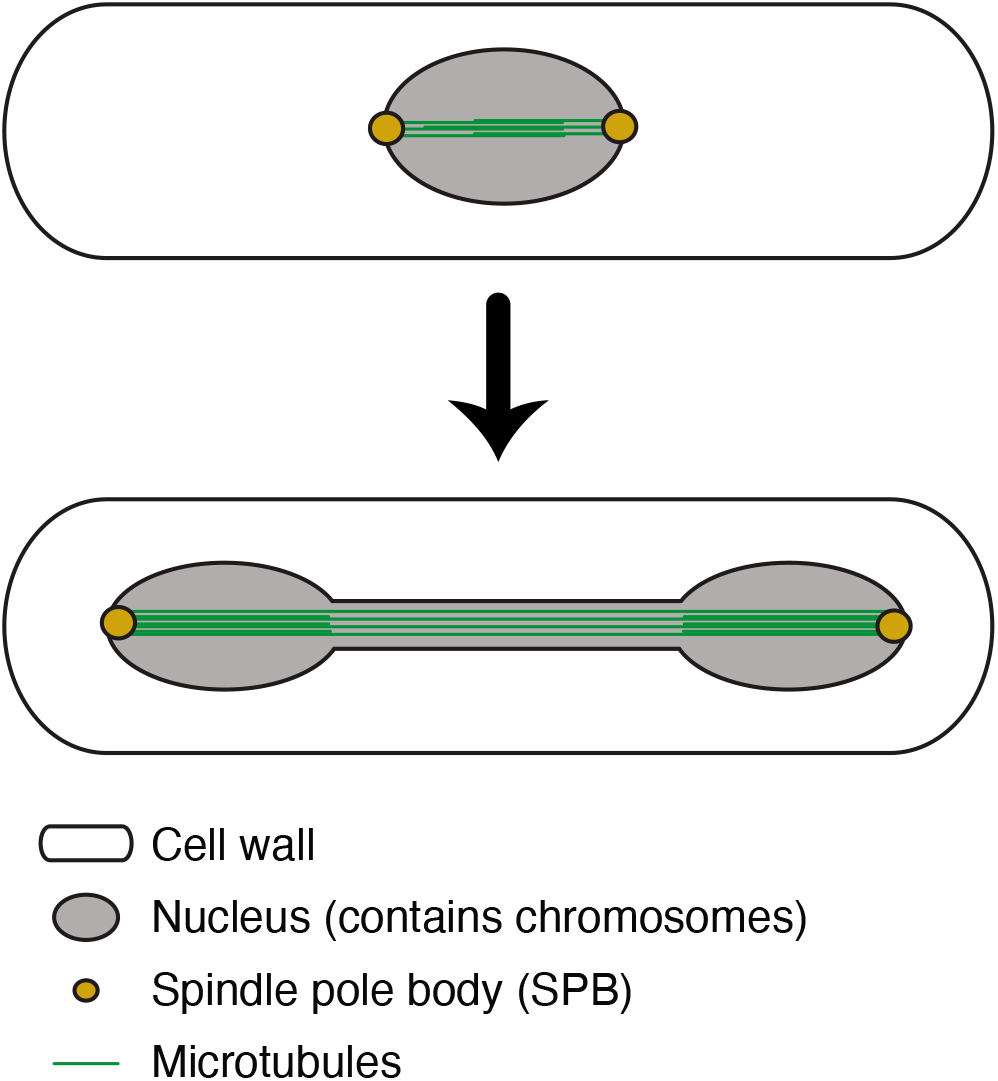
The *S. pombe* mitotic spindle, a microtubule-based cellular machine, pushes chromosomes to opposite ends of the cell during mitosis.

There are a few software packages available for specifically tracking features of *S. pombe* images. Multiple approaches are able to segment individual cells and track their growth either by brightfield imaging (37) or a fluorescent reporter (38). Notably, Li et al. (39) present an open-source software that can track *S. pombe* cell size and shape, as well as delays in metaphase/anaphase and identify abnormalities in mitosis, by interfacing with a microscope and tracking the position of individual spindle pole bodies (SPBs). However, since SPBs are diffraction-limited, their position can be quantified with techniques developed for single-molecule and super-resolution microscopy, and this approach cannot be easily extended to images of the entire spindle visualized by, for example, fluorescently-labeled tubulin.

Currently, tracking of *S. pombe* spindle elongation dynamics when visualizing the spindle itself is at best semiautomated via home-built analysis software (34,36). For example, previous work manually tracks *S. pombe* spindle lengths using kymographs (36), a semi-automated MATLAB program that requires manual segmentation of the cell (34), or an ImageJ plugin called MTrackJ (40), which requires users to manually click the desired features in each frame of the video, and records the coordinates and other information (41). Currently, to our knowledge there are no freely available, fully automated *S. pombe* spindle length tracking software packages.

We have developed a novel tool for automatically calculating the length of the mitotic spindle in *S. pombe* as the cell elongates. Using an algorithm that detects the principal axes of the spindle and then finds its ends, we have created an ImageJ plugin that tracks spindle ends and calculates the spindle length over time from any video of dividing *S. pombe* cells expressing fluorescently labeled tubulin (or other marker that labels the spindle). To assess its overall performance, we test its accuracy on synthetic microscopy images of spindles of known length at various levels of visual noise. This work presents a free, open-source software tool applicable to many areas of biological research that also serves as a baseline for future computational endeavors.

## Materials and Methods

### Yeast Strain, Growth, and Preparation for Imaging

The *S. pombe* strain used in this study was FC2012, genotype *h^+^ase1-mCherry:NatMX leu1-32::SV40-GFP-atb2 ade6-M216 leu1-32 ura4-D18 his 7^+^*, which was a gift of Fred Chang. This strain expresses both green fluorescent-tagged *α*-tubulin and mCherry-tagged Ase1p, although the latter was not imaged in these experiments. Prior to imaging, cells were grown at 25 °C on YE5S agar plates until single colonies appeared. A sample of a single colony was collected with a toothpick and inoculated in YE5S growth media. Serial dilution ranging from 1:4 to 1:16 was performed (into YE5S), and cultures were incubated overnight at 25 °C with shaking. Approximately 18 hours later, cultures in log phase were selected based on optical density. A 1 mL volume of this culture was briefly pelleted in a table-top microfuge, and the supernatant was discarded. The pellet was resuspended in 10 μL of media. An agar pad was prepared on a microscope slide, and the resuspended culture was placed on this pad and topped with a coverslip. The coverslip was then sealed with VALAP (1:1:1 Vaseline:lanolin:paraffin) and imaged immediately.

### Live cell imaging

Live-cell imaging of *S. pombe* expressing GFP-Atb2p was performed as described (42) on a Nikon Ti-E stand on an Andor Dragonfly spinning disk confocal fluorescence microscope; spinning disk dichroic Chroma ZT405/488/561/640rpc; 488 nm (50 mW) diode laser with Borealis attachment (Andor); emission filter Chroma ET525/50m; and an Andor Zyla camera. Example images shown in the text were collected with a 60x 1.49 TIRF Nikon objective. Additional images used for algorithm calibration were collected with a 100x 1.45 Ph3 Nikon objective. The algorithm worked robustly under both magnifications. Z-stacks containing 8 planes each 1 micron apart were taken to image the sample at intervals of 15 or 30 seconds (depending on the particular acquisition). Andor Fusion software was used to control the data acquisition.

### Data analysis

Before image analysis, a maximum-intensity z-projection was performed in ImageJ. Cells entering mitosis were cropped using ImageJ and saved as a TIFF stack without compression or interpolation. Linear adjustment was employed for changes in brightness and contrast. Manual tracking of the spindle length was performed with a homebuilt MATLAB program that recorded the coordinates as the user clicked on the spindle ends. Automatic tracking was performed using our spindle end-finding tool as described below. All plots were created in Python using Jupyter notebook software.

## Spindle End-Finding Algorithm Description

We have developed an algorithm, outlined in Figure 2, that can reliably detect the ends of the mitotic spindle in a microscopy image provided as a single color 16-bit TIFF file (Figure 2a). When analyzing time-lapse videos, the algorithm is performed on each frame.

**Figure 2.**
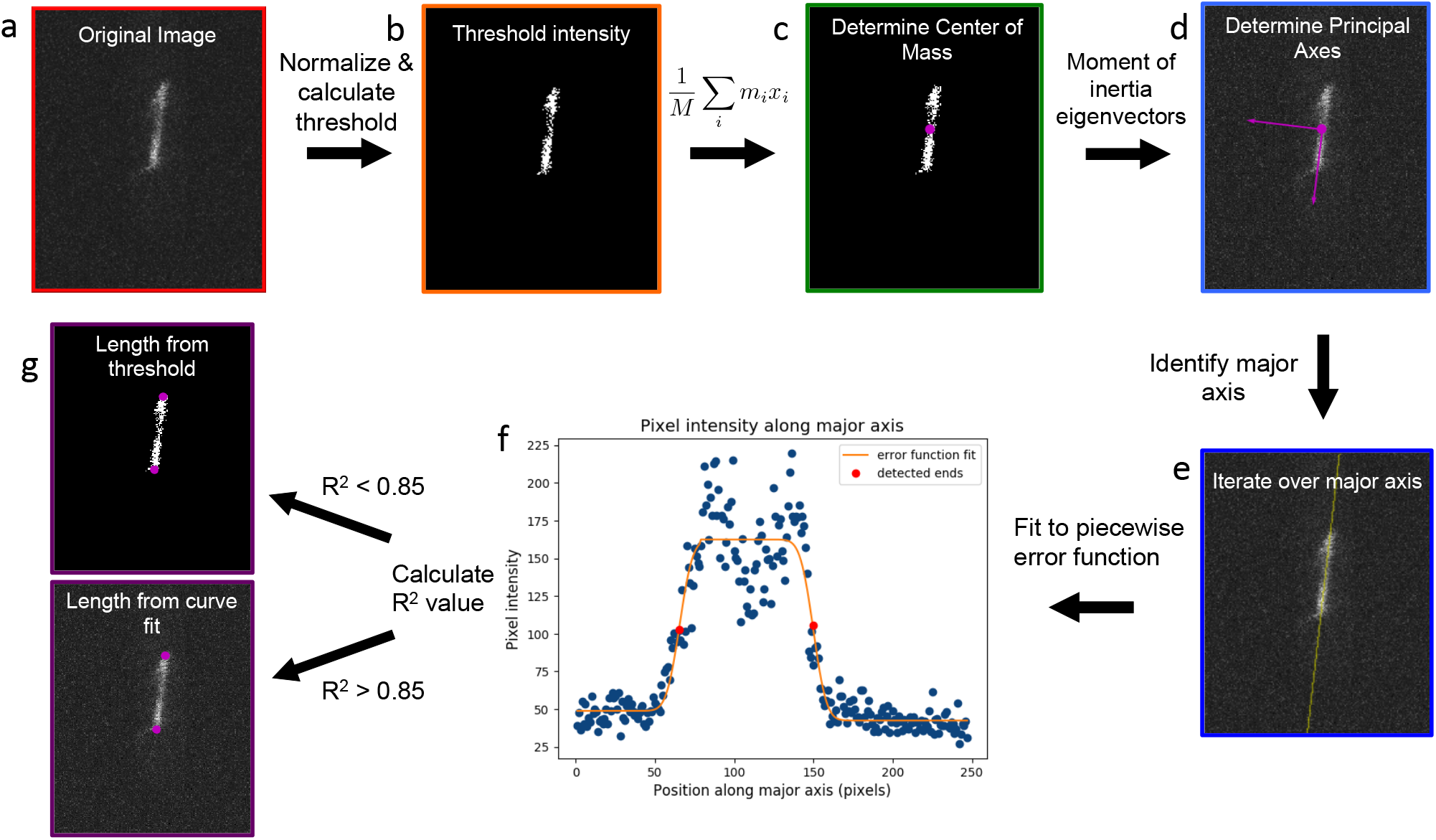
Schematic of algorithm to measure spindle length. The original image **(a)** is normalized and a threshold intensity value is interpolated and applied to the image **(b)**. The center of mass is then calculated **(c)**, and the principal axes are calculated **(d)**. We then iterate over all of the pixels along the major axis **(e)** and fit them to a spindle intensity profile function **(f)**. The length is then calculated as the distance between the ends **(g)**.

First, the following linear transformation is applied to normalize the pixel intensity values in the image to a range between 0 and 65535, the maximum pixel value for a 16-bit monochromatic image:

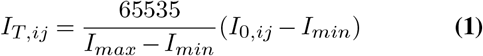

where *I_min_* and *I_max_* are the minimum and maximum pixel intensity values of the entire image, *I_T_,_i_j* represents the pixel intensity value in the transformed image, and *I_0_,_i_j* represents the pixel intensity value in the original image. The original pixel values can vary widely due to microscope settings, fluorescence background, or other factors, but it is important that the pixel intensities are all in the same range for the rest of the algorithm. This normalization ensures that the algorithm is effective for images collected under a wide range of conditions.

Next, we calculate a threshold intensity that separates the spindle from the background. Finding a suitable threshold automatically is a challenge, as it can vary with the mean intensity, the variance of the intensity, and the signal-to-noise ratio of the particular image. To overcome this challenge, we built an interpolator function that can find an appropriate threshold. To do so, we first generated a calibration dataset by recording the mean, standard deviation, skewness, and manually selected threshold intensities for 50 frames of GFP-atb2 (α-tubulin) from videos of 11 different *S. pombe* cells undergoing mitosis. We then use these as input to a linear interpolator from Scipy (43) to create a function that takes in the mean, standard deviation, and skewness of the pixel values in a given frame and returns a threshold intensity value. Pixels with intensities above this value should correspond only to the mitotic spindle. We then set all pixels with intensities below this threshold value to zero, leaving us with only the mitotic spindle in the image (Figure 2b).

To find the position and orientation of the spindle, we then calculate the center of mass coordinates of the image *(X_CM_,Y_CM_*) using pixel intensity as “mass” (Fig. 2c, Eq. 2) and the moment of inertia tensor **M** (Eq.3).

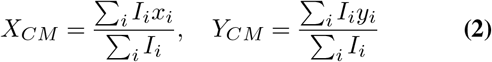

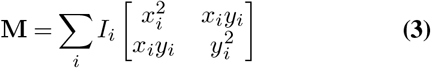

where *I_i_* is intensity for pixel *i* and *(x, y*) correspond to pixel coordinates in the image.

Next, we calculate the eigenvectors of this matrix, which correspond to the principal axes of the spindle. The vector with the larger corresponding eigenvalue is the major axis (Fig. 2d). Using this vector and beginning at the center of mass, we can then draw a straight line along the length of the spindle (Figure 2e). Beginning at one edge of the original (non-thresholded) image, we then perform a cut along this line, recording the pixel intensities along the length of the spindle (Figure 2f). In the absence of noise, this profile would look like the spindle intensity profile function shown in Figure 3 - uniformly high along the spindle and low in the background (whose level may differ at the two ends), with a steep curve at the ends of the spindle.

**Figure 3.**
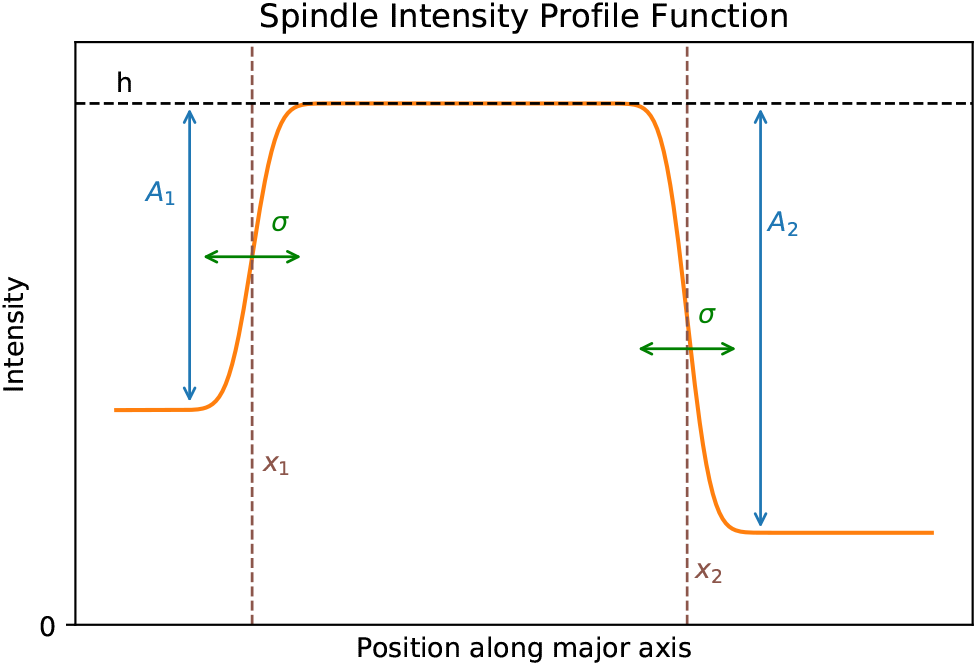
Spindle intensity profile function curve fit parameters. The curve is characterized by two error functions placed back-to-back. We find the intensity along the spindle (*h*), the vertical stretch on either end (*A*_1_ and *A*_2_), the horizontal shift on each end (*x*_1_ and *x*_2_), and the horizontal stretch for both ends (*σ*). The two error functions meet at the midpoint between *x*_1_ and *x*_2_. We calculate the location ofthe ends as the horizontal shift parameters (*x*_1_ and *x*_2_).

The shape of this curve at each end is determined by the point spread function of the microscope, but for determining the position of the spindle edge, it is well-approximated by an error function. To fit this profile and find the spindle ends, we define a function that includes two opposite-facing error functions, joined to form a piecewise, continuous function:

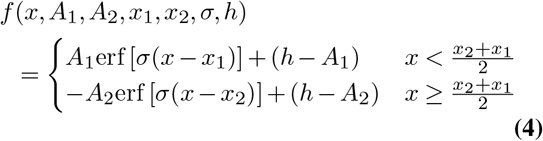

This spindle intensity profile function has 6 parameters: *A_1_* and *A*_2_, which represent the difference in intensity between the background and the spindle at each end; σ, the variance of the error function, which is determined physically by the point spread function of the microscope and corresponds approximately to the diffraction limit; *x*_1_ and *x*_2_, the positions of the beginning and end of the spindle; and *h,* the intensity of the spindle. Note that the algorithm performed best when *A*_1_ and *A*_2_ were allowed to float separately, accommodating non-uniformity in fluorescent background. While this function definition is only appropriate when |*x*_2_—*x*_1_| > σ to avoid the intersection of the two error functions, that condition will always be met for spindles with optically resolvable lengths, since σ is the width of the point-spread function of the microscope.

We then fit the intensity profile of the spindle to this function, calculating the values of the parameters defined above and shown in Figure 3. The location of the ends of the spindle are determined by the parameters *x*_1_ and *x*_2_, and we take the difference of these values to calculate the length of the spindle.

Finally, we calculate the *R*^2^ value for the curve fit. If *R*^2^ *>* 0.85 and the calculated length is less than the diagonal length of the image, the curve fits the data relatively well and we use the value of *x*_2_—*x*_1_ in Figure 3 as the spindle length. If *R*^2^ < 0.85 or the calculated length is larger than the diagonal of the image, then the calculated length is unlikely to be correct. This could occur for a number of reasons, including the signal-to-noise ratio dropping too low, or the spindle going out of focus. In this case, we return to the thresholded image, perform a cut along the major axis, and use the distance between the first and last non-zero points along that axis as the length in pixels. This is generally a less precise measurement, which is why we use it only as a backup in cases where the curve fit does not perform adequately. Having this secondary method of measuring length allows us a higher probability of continuing to accurately measure the spindle length.

This 7-step algorithm allows us to measure the length of the spindle in any given still frame of an *S. pombe* cell. For videos of cells dividing, the algorithm is repeated for each frame. The overall runtime efficiency is 𝒪(*nm*), where *n* is the number of frames and *m* is the number of pixels in each frame.

## Results

### Implementation

We have implemented the algorithm described above into a plugin for FIJI/ImageJ, a widely-used, openly available image processing software (44,45). Since ImageJ is Java-based, plugins are required to be written in Java and use the ImageJ libraries. However, much of our algorithm involves interpolation and curve fitting, and we use Python because it has readily available libraries for these functions. To allow ImageJ to run these scripts, we use system calls to run Python 3 scripts for the interpolation, curve fitting, and matrix algebra components of the algorithm. The calibration data used to calculate the threshold intensities is also available on Github, and can be adjusted by the user to fit their own microscope settings if needed. The Python scripts pass parameters to the Java program by writing them to temporary files that are then read by the main program. This requires the user’s computer to be able to run Python scripts and the necessary libraries (Scipy and NumPy), and for the file to be placed in the directory of the FIJI application, but does not require anything additional from the user after initial installation. The application was developed on MacOS but also works on Windows with additional configuration.

Once installed, the plugin can be run directly from the ImageJ plugins menu, and does not require any coding from the user. The user first opens their image as a stack within ImageJ, and then the plugin operates on the currently selected stack. After detecting the spindle ends in the image stack, the plugin outputs a comma-separated value (CSV) file of coordinates of the ends and total length of the spindle for each frame. When run, the plugin prompts the user to select a location on their device where they would like the output file to be saved. Another dialog asks the user to enter the scale of the images (in pixels/micron) if they would like the calculated lengths to be converted to microns instead of pixels.

In addition to the CSV file output, the plugin also returns a new TIFF stack of images with Regions of Interest (ROIs) placed on the detected ends (as seen in Figure 4). This allows the user to visually check the lengths determined by the algorithm, and the location of the ROIs can also be adjusted manually if needed.

**Figure 4.**
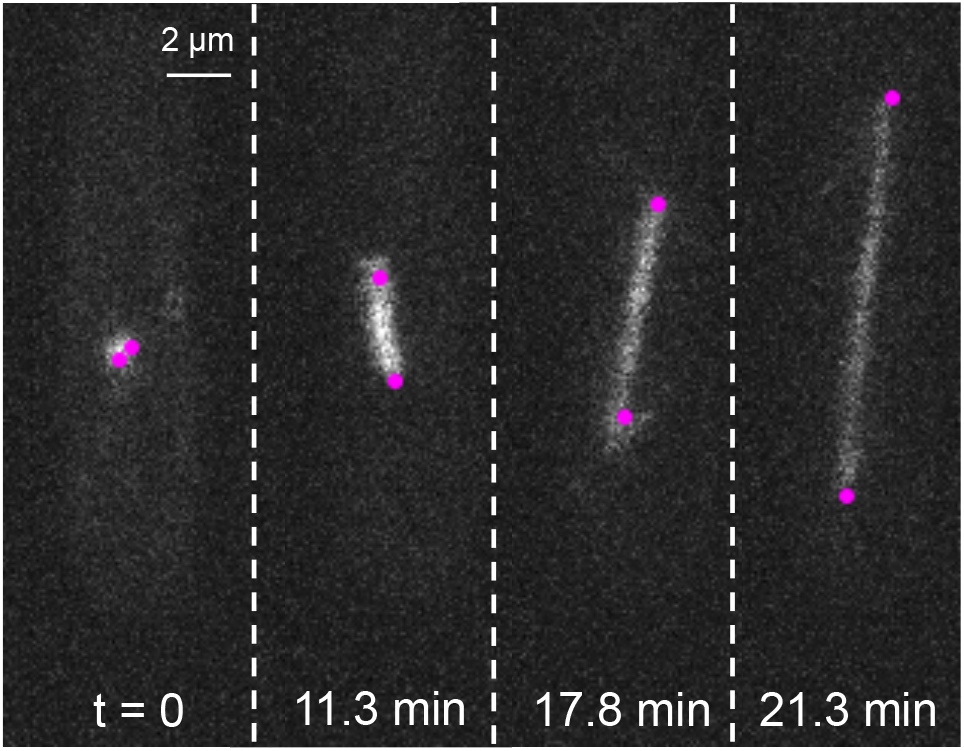
Example confocal fluorescence images of an elongating spindle in a *S. pombe* cell expressing GFP-tubulin. The ImageJ plugin places magenta ROIs at the detected ends of the spindle in each frame as it tracks the spindle’s length over time.

All of the code for this plugin is open-source and available on Github. Users can find all necessary files and installation instructions on our public GitHub repository (https://github.com/eltinglab/SpindleLengthPlugin).

### Testing with Live Cell Imaging Data

To be used effectively, it is imperative that the algorithm is comparable in accuracy for tracking the spindle as manual clicking by the user. Figure 5 compares the output of the algorithm to the lengths determined visually by a human for two different videos of *S. pombe* spindle elongation. For the majority of the points, the algorithm calculates the correct length. Most of the cases where there is a discrepancy between the length determined by the algorithm and determined visually occur towards the end of spindle elongation. This is likely because, as the spindle becomes longer, the same amount of total tubulin is spread over a longer distance, reducing the signal-to-noise ratio. Additional factors that may contribute include the spindle going out of focus, and the potential that photobleaching reduces the contrast between the spindle and the background. One way to further reduce the discrepancy between the algorithm and manual clicking is by applying a curve fit or filter to the length values produced by the algorithm. These kinds of further analyses are facilitated by the output CSV file of lengths that can easily be read in by many different kinds of software.

**Figure 5.**
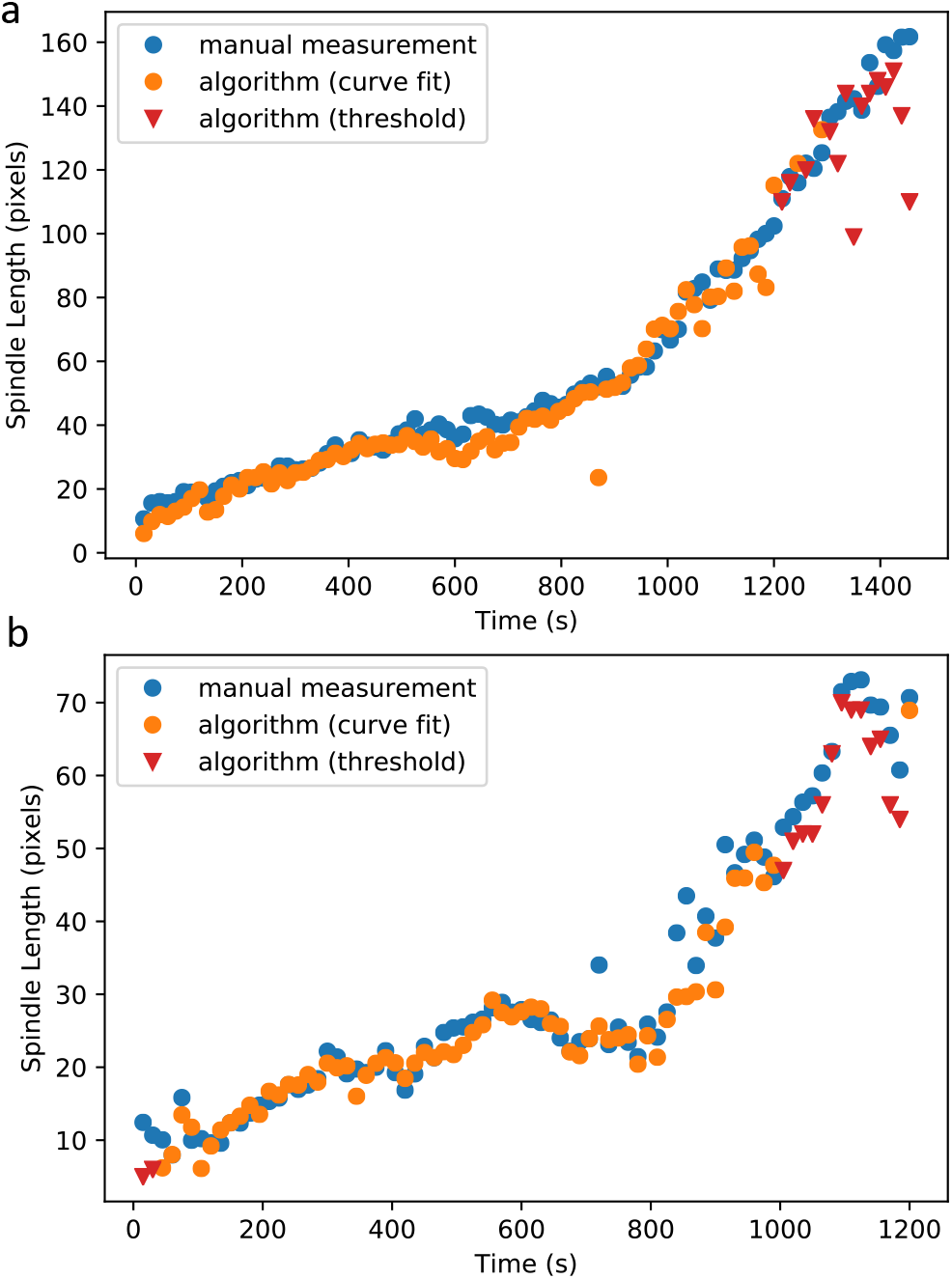
Comparison of spindle length measurements over time of the algorithm (blue) and manual clicking (orange) in two different example *S. pombe* cells (**a** and **b**). Less than 7% of frames in these videos were used in the calibration data to calculate threshold intensities. The average SNR of the cells were 6.56 and 6.55 for panels a and b, respectively. Under our imaging conditions, the scaling factor for both cell videos is 10 pixels per micron.

### Testing with Synthetic Data

To identify the limitations of the software, we created synthetic microscopy images that span a number of different conditions. To investigate how spindle length and orientation affected accuracy, we used “spindles” of 3 different lengths, and also varied the rotation angle for each of them. Additionally, we added various levels of background noise to each image to test the effect of signal-to-noise ratio (SNR). By doing this, we were able to determine the scenarios in which the algorithm performs at it best, and pinpoint factors which might weaken performance. Many of these factors can be controlled by careful choice of microscopy conditions when the images are collected, allowing the user to achieve optimal algorithm performance.

Our simulated data was 80 × 247 pixels in size, the same shape as a real recorded microscopy image that was cropped to include only one cell. To create the simulated “spindle”, a line of the desired length and angle was drawn on a black canvas 10 times as large as the final image. The line width was 12 pixels on this scaled up image, calculated based on the approximate width measured for microscopy images of real spindles. We then applied a Gaussian blur with a standard deviation of 14.7 pixels (on the scaled up image) to mimic the convolution with a point-spread function that occurs when light is diffracted through a microscope. Although the point spread function of a microscope is not strictly Gaussian, it is reasonably well-approximated by one for performing subpixel localization (46). The image was then scaled down to size, and Gaussian noise of the desired level was applied.

We explored three different spindle lengths: “long” (190 pixels), “medium” (100 pixels), and “short” (50 pixels). For each length, we tested at least three different orientation angles - the long spindle was tilted at 75, 90, and 105 degrees from the x-axis, the medium spindle at 60, 90, and 120 degrees, and the short spindle at 0, 45, 90, and 135 degrees.

For each combination of length and angle, we added five different levels of random visual noise to the image. The noise conditions are denoted by their standard deviation; the pixel intensity values are normalized to be between 0 and 65535, and the noise levels we used had standard deviations equally spaced between 0 and 20000, which correspond to a SNR range of infinity to 3.3 (lower SNR denotes more noise). Figure 6 shows the visual effects of different noise levels on the “long” spindle tilted at 75 degrees.

**Figure 6.**
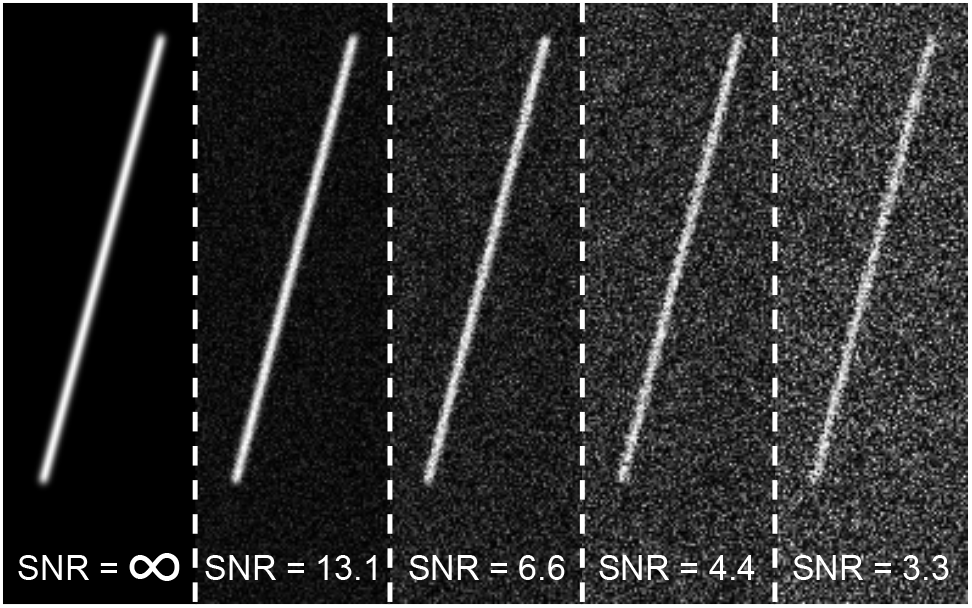
Varying levels of Gaussian noise on simulated microscopy images of a “long” spindle (190 pixels in length). Signal-to-noise ratio (SNR) is calculated as the ratio of the maximum intensity (here 65535, for a 16-bit image) to the standard deviation of the added Gaussian noise.

Each experimental condition (length/angle/noise level) was run through the plugin 15 times. Figure 7 shows some results of these simulations, and more are shown in Supplementary Figures S1, S2, and S3.

**Figure 7.**
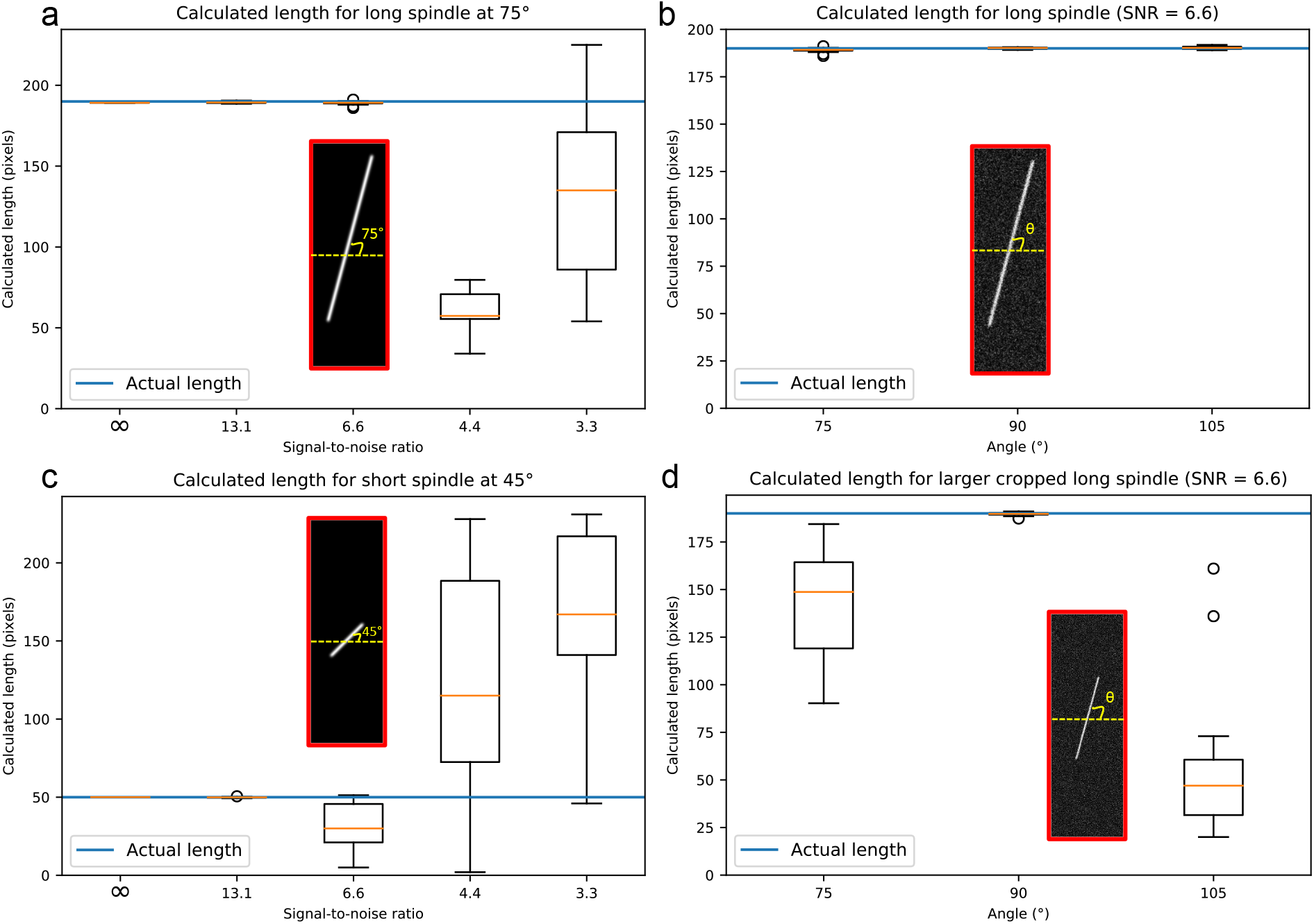
Results of simulated data. **(a)** Length vs. SNR for a long spindle (190 pixels) at an angle of 75 degrees from the x axis. **(b)** Length vs. orientation angle for a long spindle, at a constant SNR of 6.6. **(c)** Length vs. noise level for a short spindle (50 pixels) at a 45 degree angle. **(d)** Length vs. angle for a long spindle in a frame twice as big (160 × 494), at constant SNR of 6.6. Insets show the synthetic spindle with the angles denoted in yellow. All images are 80 × 247 except in (d). The blue line indicates the actual spindle length. There were 15 trials for each experimental condition.

We first assessed the performance of the algorithm over a range of noise levels where spindle length and angle were held constant, for a long spindle moderately aligned with the long axis of the image (7a). The algorithm performs well, producing lengths close of the actual length (denoted with the blue line), until a SNR of 4.4, where it starts severely underestimating the spindle length. This likely denotes the level of visual noise that starts significantly affecting the algorithm’s performance. If there is too much visual noise, the principal axes calculations can be thrown off, and as a result, the spindle will not be detected correctly. An additional possibility is that, even if the major axis is correctly identified, the level of noise makes it impossible for the spindle intensity profile function to fit the data well. Evidence for both of these scenarios was found in the image results of these simulations.

We next fixed the SNR to 6.6, the level where the algorithm began to underperform (Figure 7a), and assessed perofrmance as a function of spindle orientation angle (Figure 7b). When the spindle is aligned either along the long axis of the image, the algorithm performs best. However, we note that at at higher noise, the spindle is more likely to be correctly detected if it is straighter (seen in Figure S1).

To assess performance for short spindles, we repeated the simulation from Figure 7a on a short spindle tilted at 45 degrees (Figure 7c). At low noise levels, the algorithm performs well, but begins to diverge from the actual value at an SNR of 6.6. This is a somewhat higher than the SNR at which the algorithm performance worsens for long spindles. This difference is likely due to the number of total pixels in the image that are in the spindle. If there are fewer of them, as when there is a smaller spindle, it takes less visual noise to dominate the image. At higher noise levels, the algorithm performs similarly poorly for all lengths, but is more likely to underestimate the lengths of long spindles and to overestimate the lengths of short spindles.

Finally, we tested if the crop size of the image affects the algorithm’s performance by repeating the simulation from Figure 7b but with an image canvas sized to 160 × 494, twice as large as the previous images (Figure 7d). In this case, the algorithm performs quite poorly at low SNRs, vastly overestimating the spindle length with margins of error much higher than in the more closely cropped image. This likely occurs for the same reason as with the short spindle at higher noise levels: since the spindle takes up less of the whole image, it is more easily overwhelmed by noise. However, this issue is also resolved at higher SNRs. The noise tolerance of our algorithm is similar to image analysis software used for superresolution localization, which has been described as failing with SNR < 5 (12).

We also show additional examples of results of these simulations for long (Figure S1), medium (Figure S2), and short (Figure S3) spindles under a variety of conditions. The results show that overall, the algorithm works well across all lengths and angles when there the visual noise is kept to a manageable level ⪆ 6, which is readily achievable with standard fluorescence microscopy, as shown in Figure 4. When the noise becomes more prominent, performance worsens, especially for shorter or more angled spindles. In order to get the best results with the FIJI plugin, users should try to keep visual noise in the microscopy image to a minimum, and also to crop the video as close to the desired cell as possible. Additionally, a higher image resolution is conducive to more accurate results.

## Conclusions

We present a new computational tool that automatically measures *S. pombe* spindle length over time. Using an algorithm that calculates the center of mass and principal axes of a still frame of a mitotic spindle, we can accurately find the ends of the spindle and calculate its length. We have implemented this algorithm into a FIJI/ImageJ plugin that can be run on TIFF stacks and does not require any coding. The plugin outputs a CSV file of the spindle length in each frame that can be used for further analysis in much less time than it would take to manually measure the spindle length over time.

To analyze the effectiveness of the software, we created synthetic microscopy images of mitotic spindles of various lengths and angles, and added different levels of visual noise to each case. The results of these simulations showed that at reasonable noise levels, the plugin works well across all lengths and angles within the image. At lower signal-to-noise levels, the software performs better on longer spindles rather than shorter ones, and on spindles aligned with the image axis over angled ones. Additionally, the algorithm yields better results when the video is closely cropped to a single cell, rather than zoomed out.

This algorithm should be immediately useful and accessible for users who wish to automate the quantification of *S. pombe* spindle dynamics with minimal input from the user and without the necessity of programming skills. This application would be particularly useful for cases in which quickly quantifying the behavior of many cells is important, such as for conducting a mutagenic screen assessing mitotic progression. The current plugin can also serve as a basis to measure additional spindle features. For example, orientation angle or pole body location could easily be quantified with the current algorithm. This approach could also be expanded to other cell biological systems, such as spindles of other organisms or for tracking cytoskeletal filament ends outside the context of the spindle or *in vitro.*

All of the code for this plugin, as well as installation instructions, are available on our public GitHub repository (https://github.com/eltinglab/SpindleLengthPlugin).

## ACKNOWLEDGMENTS

The authors thank Eva Johannes and Mariusz Zareba of the NC State Cellular and Molecular Imaging Facility, Caroline Laplante, Arthur Molines, Fred Chang, Natalie Chazal, and members of the Elting lab for helpful advice and discussion. A.S.M.U acknowledges support by a NC State Park Scholarship and a Goldwater Scholarship. M.W.E acknowledges support by NIH 1R35GM138083. A.F.K. acknowledges support by the National Science Foundation under grant DMR-1752713.

## Supplementary Figures

**Figure S1.**
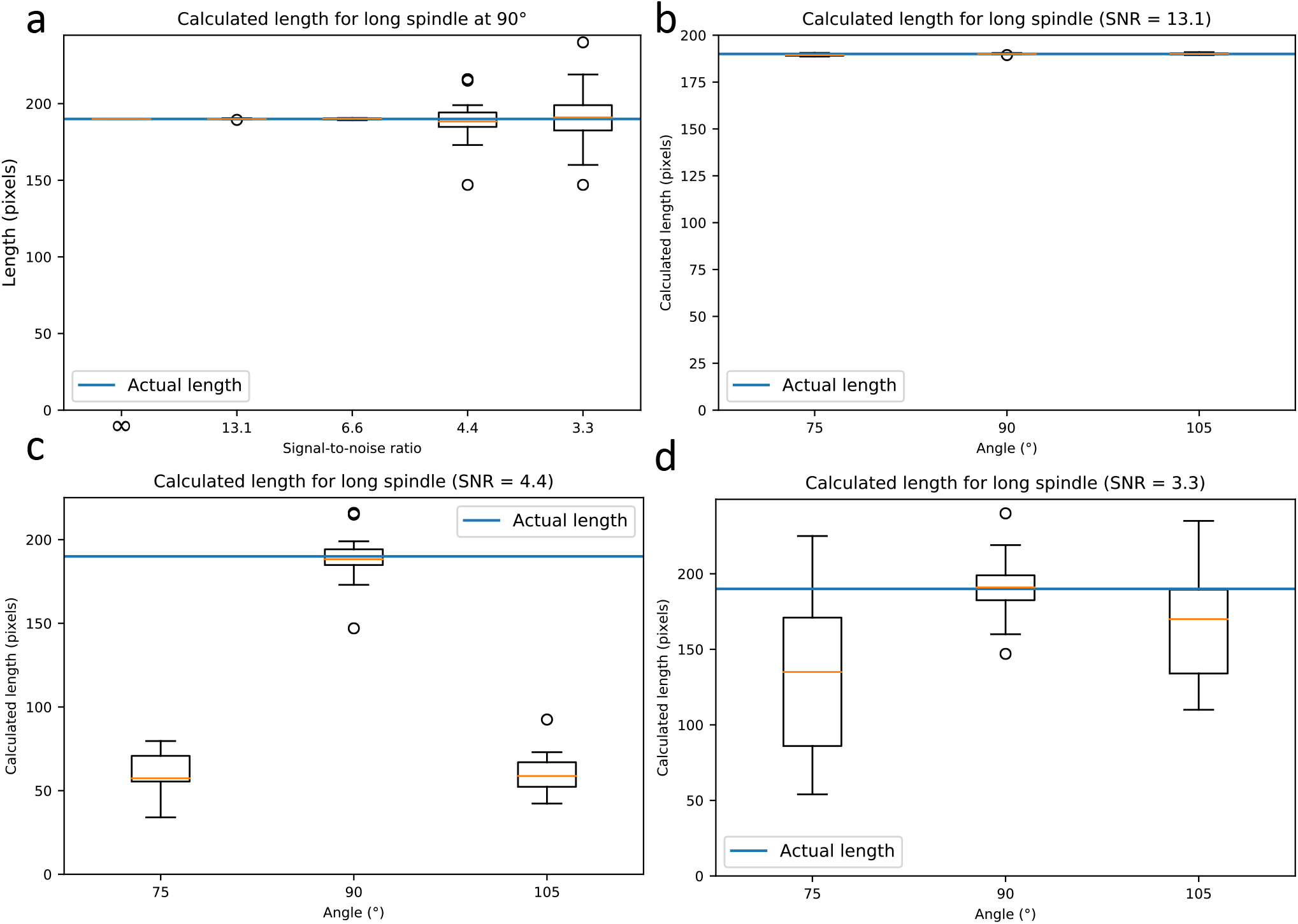
Additional results of simulated data analysis for the “long” spindle (190 pixels). **(a)** Length vs. SNR for a long spindle at an angle of 90 degrees from the x axis. **(b)** Length vs. orientation angle for a long spindle, at a constant SNR of 13.1. **(c)** Length vs. orientation angle for a long spindle, at a constant SNR of 4.4. **(d)** Length vs. orientation angle for a long spindle, at a constant SNR of 3.3. All images are 80 × 247. The blue line indicates the actual spindle length. There were 15 trials for each experimental condition.

**Figure S2.**
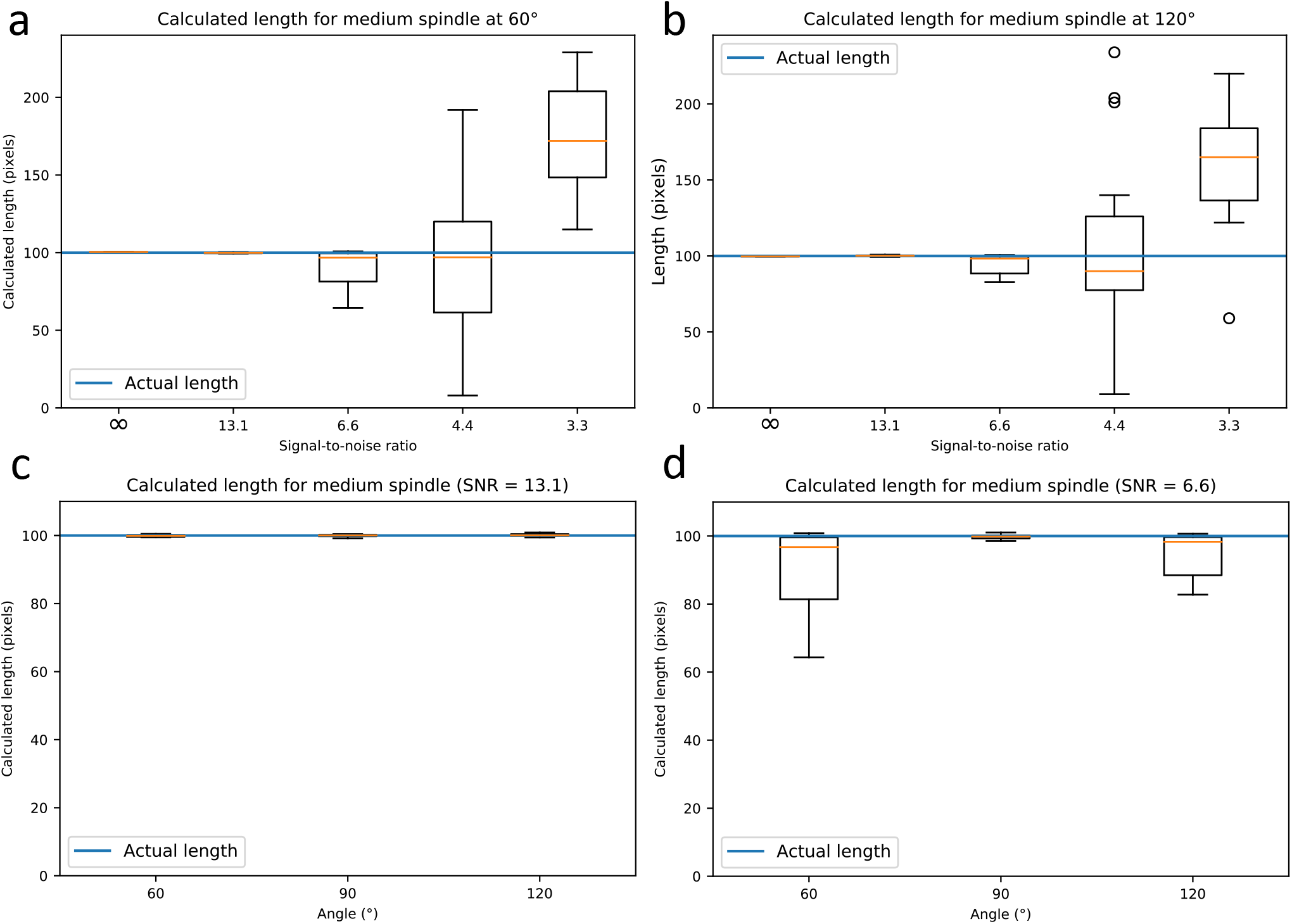
Additional results of simulated data analysis for the “medium” spindle (100 pixels). **(a)** Length vs. SNR for a medium spindle at an angle of 60 degrees from the x axis. **(b)** Length vs. SNR for a medium spindle at an angle of 120 degrees from the x axis. **(c)** Length vs. orientation angle for a medium spindle, at a constant SNR of 13.1. **(d)** Length vs. orientation angle for a medium spindle, at a constant SNR of 6.6. All images are 80 × 247. The blue line indicates the actual spindle length. There were 15 trials for each experimental condition.

**Figure S3.**
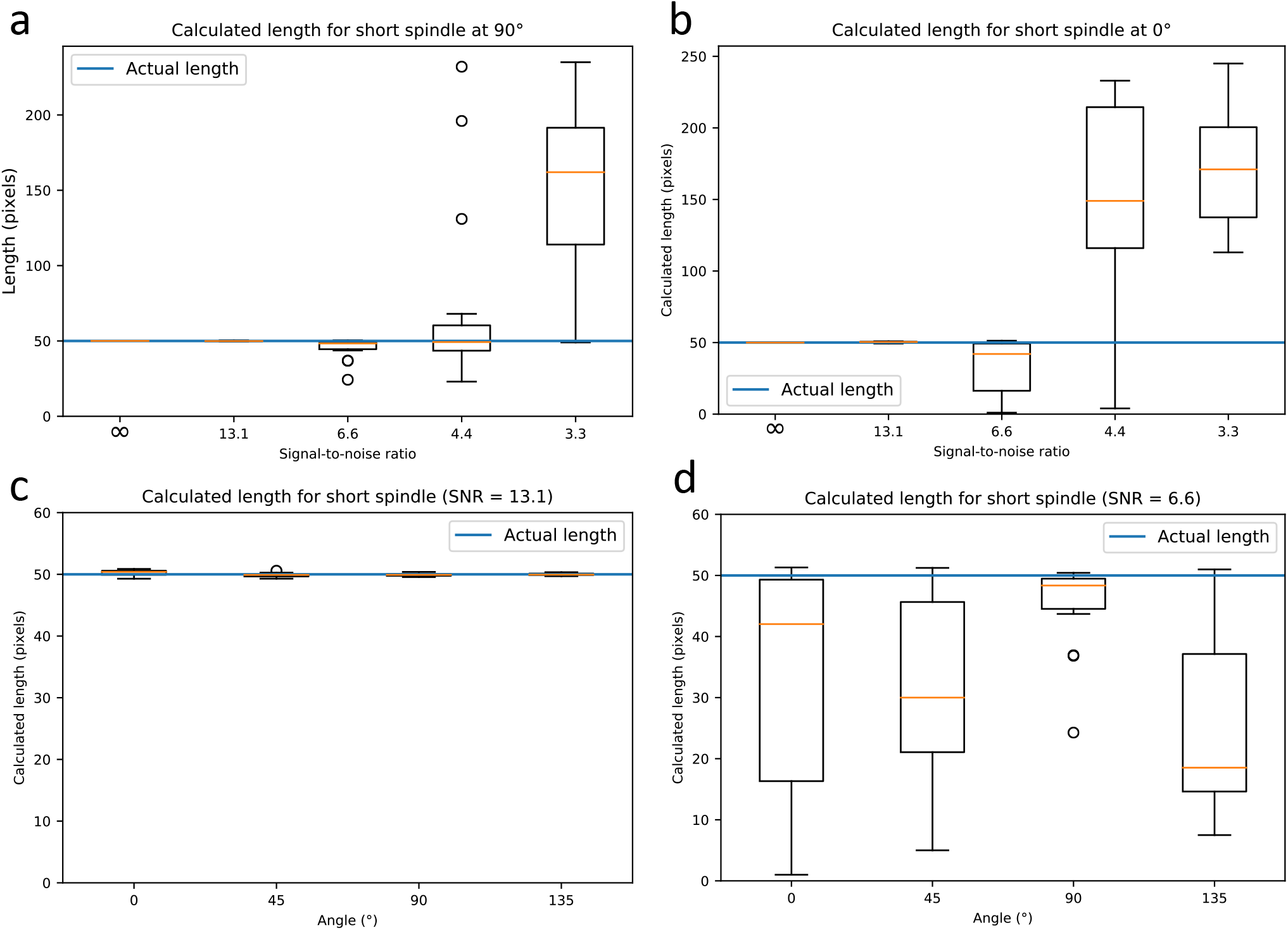
Additional results of simulated data analysis for the “short” spindle (50 pixels). **(a)** Length vs. SNR for a short spindle at an angle of 90 degrees from the x axis. **(b)** Length vs. SNR for a short spindle at an angle of 0 degrees from the x axis. **(c)** Length vs. orientation angle for a short spindle, at a constant SNR of 13.1 **(d)** Length vs. orientation angle for a short spindle, at a constant SNR of 6.6. All images are 80 × 247. The blue line indicates the actual spindle length. There were 15 trials for each experimental condition.

